# Selective Auditory Attention Decoding with a Two-Node Wireless EEG Sensor Network

**DOI:** 10.64898/2026.02.17.706305

**Authors:** Simon Geirnaert, Ruochen Ding, Alexander Bertrand

**Author notes:** Author to whom any correspondence should be addressed. **E-mail:**.

## Abstract

**Objective:** Selective auditory attention decoding (sAAD) enables neuro-steered hearing devices by identifying the attended speaker in a multi-speaker environment from neural activity recorded with electroencephalography (EEG). Despite algorithmic progress, practical deployment remains constrained by a lack of wearable, unobtrusive, and fully wireless EEG acquisition solutions. We therefore evaluate whether reliable sAAD can be achieved under the realistic hardware constraints of a wireless EEG sensor network (WESN) of miniaturized, galvanically isolated EEG sensor nodes.

**Approach:** We use a WESN of two synchronized, compact around-ear EEG sensor nodes worn bilaterally. Each node provides four local EEG channels derived from five pre-gelled electrodes, including a local reference. Sample-wise wireless synchronization of data from both nodes enables joint processing as an eight-channel EEG. On a newly recorded dataset, we apply correlation-based stimulus decoding, followed by hidden Markov model (HMM)-based post-processing.

**Main results:** Stimulus decoding achieves an average sAAD accuracy of 69.24% on 60 s decision windows, comparable to previously reported wired around-ear EEG systems that measure long-distance scalp potentials. HMM-based post-processing further improves this to a steady-state accuracy of 77.17% with an average simulated attention switch detection time of 32.79 s. Combining sensor nodes at both ears outperforms single-ear configurations, primarily through redundancy rather than complementary spatial information. Finally, a fixed bipolar configuration using four electrodes per ear, yielding three channels, suffices to maintain performance.

**Significance:** These results demonstrate the practical feasibility of sAAD using a fully wireless, galvanically isolated around-ear WESN and establish a realistic performance benchmark under practical hardware constraints.

## 1 Introduction

Selective auditory attention to one out of multiple competing speech streams can be decoded from neural responses, for example, recorded using electroencephalography (EEG) (Ding and Simon, 2012; Mesgarani and Chang, 2012; O’Sullivan et al., 2014). This capability enables the development of neuro-steered hearing devices, such as hearing aids and cochlear implants, in which the user can steer the device towards the speech stream of interest through attention-driven speech enhancement (Van Eyndhoven et al., 2017; Geirnaert et al., 2021c). Such neuro-steered hearing devices therefore hold significant potential to assist hearing-impaired listeners in challenging cocktail party scenarios. However, realizing this potential in practice requires balancing decoding performance with strict constraints on wearability, comfort, and long-term usability imposed by hearing device form factors.

In recent years, substantial progress has been made in multiple components required for neurosteered hearing devices. From a methodological perspective, selective auditory attention decoding (sAAD) algorithms have been further developed, for example, by employing deep neural networks (Nguyen et al., 2025; Sridhar et al., 2025), by introducing advanced post-processing strategies for correlation-based neural tracking methods (Miran et al., 2018; Heintz et al., 2025), or by developing unsupervised variants (Geirnaert et al., 2021b, 2022). In parallel, the integration of sAAD algorithms with speech separation and enhancement pipelines has been investigated (Van Eyndhoven et al., 2017; Ceolini et al., 2020), as well as validation in more realistic acoustic environments (Choudhari et al., 2024; Van de Ryck et al., 2025) and in hearing-impaired populations (Fuglsang et al., 2020). Despite these advances, a major - if not the most important one - remaining obstacle to practical deployment is the integration of wearable, unobtrusive neuromonitoring sensors with sAAD algorithms. In this work, we focus on this bottleneck.

EEG is commonly considered the preferred neuromonitoring modality for neuro-steered hearing devices because of its excellent temporal resolution, which is essential for neural tracking-based sAAD, and its non-invasive, relatively low-cost, and wearable nature (Nicolas-Alonso and Gomez-Gil, 2012). These characteristics make EEG particularly attractive for integration into everyday hearing devices. However, the vast majority of sAAD algorithms have been developed and validated using high-density, full-scalp wet EEG systems, which are impractical for long-term, real-world use due to their bulkiness and obtrusiveness. Enabling sAAD in hearing devices therefore necessitates wearable, unobtrusive, and ideally dry EEG solutions.

One promising approach towards unobtrusive EEG monitoring is the use of a *wireless EEG sensor network* (WESN), consisting of multiple miniaturized EEG sensor nodes (mini-EEGs) that record EEG locally at different scalp locations (Bertrand, 2015). A defining characteristic of such WESNs is the absence of a shared electrical reference through a wired connection between sensor nodes, as the mini-EEG nodes are galvanically isolated from each other by design. As a direct consequence, each node can only record short-distance, within-node scalp potentials, in contrast to conventional EEG systems that measure long-distance potentials referenced across the scalp. This limitation introduced by miniaturization can be mitigated by deploying multiple mini-EEGs in a network, thereby providing a spatially diversified representation of neural activity that is required for effective sAAD (Geirnaert et al., 2025). At the same time, the absence of wired connections reduces susceptibility to electromagnetic interference, minimizes motion-induced wire artifacts, and increases robustness to individual node failures (Bertrand, 2015; Ding et al., 2025b).

Assuming idealized mini-EEGs that can be flexibly placed on the scalp, the optimal sensor locations for sAAD have been investigated in a data-driven manner for correlation-based stimulus decoding algorithms (Mirkovic et al., 2015; Mundanad Narayanan and Bertrand, 2020). Mundanad Narayanan and Bertrand (2020) demonstrated that, starting from a full-scalp EEG configuration, the number of channels could be reduced from 64 to 10 using a greedy utility-based selection algorithm without loss in sAAD performance, achieving approximately 85–90% accuracy on 60 s decision windows. However, this analysis did not account for the constraint of short inter-electrode distances inherent to mini-EEGs. To address this, Mundanad Narayanan et al. (2021) emulated mini-EEGs using ultra-high-density 255-channel EEG recordings and showed that sAAD performance remained stable for inter-electrode distances down to 3 cm, with optimal sensor locations situated above the temporal lobe, near the auditory cortex, as expected for auditory decoding. Additionally, Mundanad Narayanan and Bertrand (2020) demonstrated through emulation the potential benefit of using two-channel rather than single-channel mini-EEG sensor nodes. The addition of a second channel allows each node to record EEG potentials in two orthogonal directions, thereby allowing to capture source dipoles in any direction.

Importantly, the aforementioned studies assume idealized sensor placement. In practice, sensor locations are constrained by form-factor considerations, user comfort, and presence or absence of hair. Consequently, practical mini-EEG implementations for sAAD have primarily focused on flex-printed arrays placed around and behind the ear (e.g., cEEGrid) (Mirkovic et al., 2016; Holtze et al., 2022; Nogueira et al., 2019; Zhu et al., 2025; Geirnaert et al., 2025) and customized in-ear EEG devices (Fiedler et al., 2017; Geirnaert et al., 2025). Using correlation-based stimulus decoding, which relies on neural tracking, typical decoding accuracies of approximately 70% for wet around-ear EEG (Mirkovic et al., 2016; Holtze et al., 2022; Geirnaert et al., 2025) and 60% for dry in-ear EEG have been reported on 60 s decision windows (Geirnaert et al., 2025). Furthermore, Geirnaert et al. (2025) emulated galvanic isolation between the ears by applying same-ear rereferencing, demonstrating no performance loss for around-ear EEG, but a 4.1 percentage point decrease in accuracy for in-ear EEG. They further showed that combining ear-centered mini-EEGs with a small number of scalp electrodes can yield near full-scalp performance, for example by adding only three scalp electrodes to in-ear EEG.

Nevertheless, in all prior work on mini-EEGs and WESNs for sAAD, the sensor nodes were still physically connected by wires, and the effects of miniaturization resulting in true galvanic isolation and exclusive reliance on short-distance potentials have only been studied through emulation. In this work, we address this gap by employing a practical realization of a WESN for sAAD using two fully galvanically isolated around-ear sensor nodes. Specifically, we build on the EEG-Linx platform introduced by Ding et al. (2025b), using two compact, synchronized wireless EEG sensor nodes with a printed circuit board (PCB)-size of 2 × 3 cm, each equipped with an embedded processor and wireless radio and each providing four EEG channels. Crucially, no wired connection is present between the two ears, as each node operates independently. Ding et al. (2025a) previously demonstrated the feasibility of this setup for decoding auditory steady-state responses (ASSRs), showing improved robustness. Here, we extend this platform to sAAD, providing a proof of concept of a two-node, four-channel-per-node WESN for practical neuro-steered hearing devices and enabling an empirical investigation of the effects of miniaturization. We use a correlation-based stimulus decoding algorithm rather than alternative direct classification approaches that decode, for instance, the spatial focus of attention (Geirnaert et al., 2021a; Vandecappelle et al., 2021). Although such approaches can achieve higher accuracies at shorter decision windows, they are also much more susceptible to overfitting, shortcut learning, and overly optimistic performance estimates (Puffay et al., 2023; Rotaru et al., 2024; Ivucic et al., 2024; Yan et al., 2025). Moreover, we show that the accuracy can be substantially improved by incorporating the recently proposed hidden Markov model post-processing approach for correlation-based stimulus decoding (Heintz et al., 2025), thereby establishing a state-of-the-art sAAD performance benchmark under the realistic hardware constraints of neuro-steered hearing devices.

In Section 2, we describe the employed sensor nodes, the EEG-Linx platform, and the recorded dataset. The sAAD algorithm, preprocessing, and evaluation procedure are detailed in Section 3. The results are reported in Section 4 and discussed in Section 5.

## 2 Methods: Data Collection

This section describes the around-ear WESN platform employed for EEG data collection in Section 2.1 and the recorded dataset in Section 2.2.

### 2.1 Wireless EEG Sensor Nodes for the WESN

EEG was recorded using two battery-powered around-ear sensor nodes worn bilaterally (one per ear; see Figure 1), which streamed data wirelessly to a hub node that was connected to an acquisition PC via USB. Each node recorded four EEG channels in a (local) common-reference montage, with four around-ear electrodes referenced to a single local electrode on the same side. This resulted in eight EEG channels in total across both ears. Since the two ear-worn nodes were galvanically isolated, referencing was necessarily performed independently per ear, restricting measurements to short-distance, same-ear potentials. The system operated without an extra driven right leg (DRL) or ground electrode, as input biasing was provided by the analog front-end circuitry on each node (Ding et al., 2025b). The same WESN setup has been successfully used to record ASSRs in Ding et al. (2025a), providing initial evidence for the validity of the setup. More detailed information on the WESN than below can be found in Ding et al. (2025b,a).

**Figure 1.**
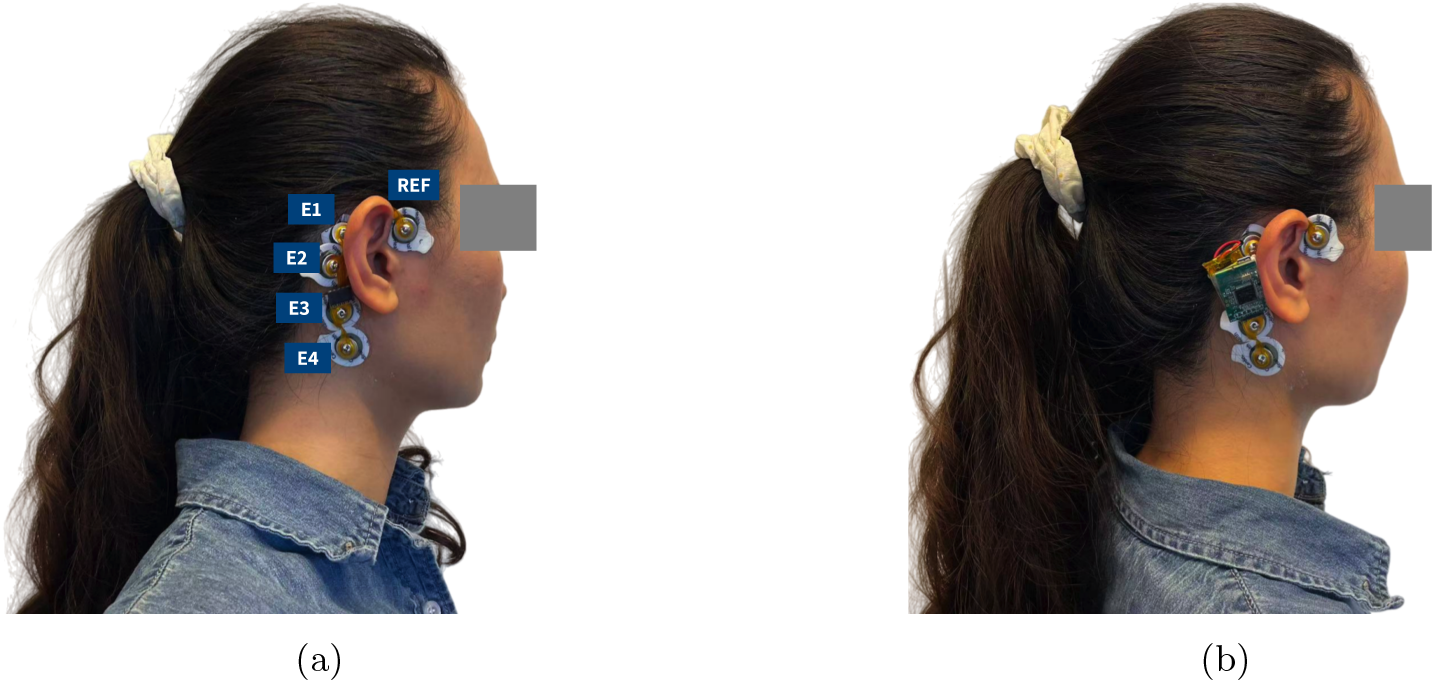
**(a)** The reusable C-shaped flex-PCB carrier with disposable snap-type pre-gelled ECG electrodes. Recording electrodes are labeled E1–E4, and the local reference electrode (REF) is placed in the preauricular region anterior to the tragus at the temporal hairline. **(b)** An example of the EEG-Linx sensor node worn behind the ear. (Photograph is of one of the authors, who has provided consent for inclusion.)

#### 2.1.1 Wireless EEG sensor nodes

The wearable EEG sensor node, originally introduced and validated in Ding et al. (2025b), is realized on a compact double-sided, six-layer PCB (approximately 2× 3 cm). Each node integrates (i) an EEG analog front-end, (ii) an embedded processing and wireless communication unit, and (iii) power-management circuitry. The analog front-end provides the interface to the around-ear electrodes and applies on-node biasing to set the input operating point during recordings. EEG signals were sampled at 250 Hz and streamed wirelessly to the hub node during the experiments. Each node was powered by a 3.7 V, 250 mAh Li-Po battery.

#### 2.1.2 Electrode interface and around-ear electrode layout

To facilitate rapid preparation and ensure hygiene across recording sessions, the electrode interface was implemented using disposable snap-type ECG electrodes mounted on a reusable flexible PCB carrier. The carrier consisted of a C-shaped flex PCB with a grid layout following the peri-auricular contour, similar to the cEEGrid configuration (Bleichner and Debener, 2017). Integrated snap studs provided both the mechanical and electrical interface between the flex PCB and the electrode pads, allowing electrodes to be attached and replaced without tools.

Commercial foam ECG electrodes (Kendall Arbo electrodes H124SG) with a diameter of 24 mm and pre-applied conductive hydrogel were used as disposable skin-contact elements. No additional gel application was required. Prior to electrode placement, the skin was cleaned using a dedicated skin-preparation gel. The disposable electrodes were then snapped onto the flex PCB carrier, after which the carrier was positioned around the ear according to the intended montage.

Electrode positions along the carrier were labeled E1–E4 following their order from superior-anterior to inferior-posterior around the pinna, as shown in Figure 1a. The local reference electrode (REF) was placed in the preauricular region, anterior to the tragus at the temporal hairline. After each recording session, the disposable ECG electrodes were detached from the snap studs and discarded, while the flex PCB carrier was retained for reuse after routine cleaning.

#### 2.1.3 Wireless acquisition, synchronization, and logging with EEG-Linx

During the experiments, the ear-worn sensor nodes transmitted EEG data wirelessly to a hub node connected to the acquisition computer via USB (Ding et al., 2025b). Since each node samples EEG using its own local oscillator, the data streams exhibit relative clock offsets and drift. EEG-Linx compensates for these effects using packet timestamps and clock-drift compensation to align the streams in time according to the clock of the hub node, which acts as the universal time for the WESN nodes. In parallel, the stimulus presentation computer transmitted trigger events to the hub node during audio playback. These triggers were recorded and time-stamped by the hub node, allowing to align the EEG data with the stimulus timeline and to segment trials in the subsequent analysis.

EEG-Linx was used to configure the sensor nodes, monitor signal quality during experimental setup, and log the synchronized EEG streams together with corresponding stimulus trigger events (Ding et al., 2025b).

### 2.2 Dataset

This section describes a newly recorded sAAD dataset acquired using the wireless two-node around-ear EEG setup, as described in Section 2.1. The experimental protocol was approved by the KU Leuven Social and Societal Ethics Committee. The complete dataset is made publicly available (Ding et al., 2026).

#### 2.2.1 Participants

Sixteen adults (seven women, nine men) between 20 and 51 years of age (mean: 29.8, standard deviation: 8.4), all with self-reported normal hearing, participated in the study.

All participants were proficient English speakers (non-native, except for one) and provided written informed consent prior to the experiment.

#### 2.2.2 Protocol and stimuli

During the selective auditory attention experiment, participants were instructed to attend to one of two competing speech streams while seated in a regular room without electrical shielding. The two speech streams consisted of stories read by English-speaking male narrators (taken from https://openslr.org) and were presented simultaneously through two loud-speakers positioned at approximately *±* 30^*°*^ azimuth and at a distance of 1 m from the participant.

The sound level was adjusted individually to a comfortable listening level. No additional manipulations were applied to the auditory stimuli.

The stories were divided into ten parts, resulting in an experimental protocol comprising ten trials of 5 min each, with a maximum total duration of 50 min. Prior to each trial, the experimenter instructed the participant to attend to either the left or right speaker, with the attended side alternating between successive trials. Participants were allowed to take breaks between trials at their own discretion. No explicit behavioral attention assessment (e.g., comprehension questions) was performed.

For various reasons (depleted battery, premature ending of the experiment at the participant’s request, (human) errors in the configuration before each trial, and other technical issues), the number of trials and duration of individual trials varied slightly across participants, sometimes leading to incomplete or missing trials. Approximately 80% of all 5 min trials across the 16 participants were completed in full, resulting in an average recording duration of 42 min per participant (standard deviation: 6.7 min; minimum duration: 25 min). All trials, including incomplete ones, were retained and included in the analysis as all participants had a sufficient number of trials.

## 3 Methods: Data Analysis

Section 3.1 presents the correlation-based linear stimulus decoding algorithm for sAAD, complemented by a hidden Markov model (HMM) post-processor (Heintz et al., 2025). Preprocessing steps for the EEG and speech signals are detailed in Section 3.2, and the evaluation procedure is described in Section 3.3.

### 3.1 Selective auditory attention decoding algorithm

The employed sAAD framework consists of two components:

1. A correlation-based neural tracking algorithm using linear stimulus decoding to generate attentional scores for each competing speech stream (Section 3.1.1).
2. An HMM-based post-processor that exploits temporal regularities in attentional processes by integrating neural tracking correlations across successive decision windows (Section 3.1.2).

#### 3.1.1 Correlation-based stimulus decoding

We employ a linear stimulus decoding approach for sAAD (O’Sullivan et al., 2014; Biesmans et al., 2017; Geirnaert et al., 2021c), which is widely regarded as one of the most robust methods and has been successfully applied in combination with around-ear EEG (Mirkovic et al., 2016; Holtze et al., 2022; Geirnaert et al., 2025). The method relies on neural tracking, whereby neural responses synchronize more strongly over time with the attended speech signal than with the unattended one (Ding and Simon, 2012; Mesgarani and Chang, 2012; O’Sullivan et al., 2014). A linear decoder is trained to reconstruct a representation of the attended speech signal from the EEG responses, after which the reconstructed signal is correlated with candidate speech representations to infer the attended speaker.

As a speech representation, the popular acoustic envelope is used (O’Sullivan et al., 2014; Biesmans et al., 2017; Geirnaert et al., 2021c). Below, a brief summary of the decoding approach is provided; extensive explanations are available in the literature (O’Sullivan et al., 2014; Biesmans et al., 2017; Geirnaert et al., 2021c, 2022).

Given a training set of *T* samples of time-lagged EEG data **X***∈* ℝ^*T ×CL*^ (where *C* denotes the number of EEG channels and *L* the number of time lags) and the attended speech envelope **y**_att_ *∈* ℝ^*T*^, a spatio-temporal linear decoder **d** *∈* ℝ^*CL*^ is learned to reconstruct the attended envelope **ŷ** = **Xd** by solving

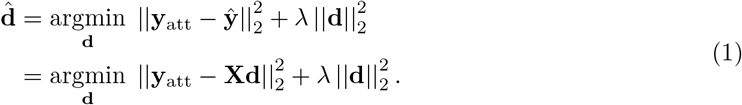

Here, **X** is a block Hankel matrix containing lagged EEG samples per channel. The time lags are chosen acausally, spanning the interval [0, 400]ms to capture neural responses following the stimulus (Geirnaert et al., 2022). This least-squares formulation is equivalent to maximizing the Pearson correlation between the reconstructed and attended speech envelopes (Biesmans et al., 2017). The closed-form solution of Eq. (1) is given by

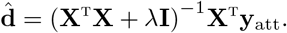

The *L*_2_-norm regularization parameter *λ* is determined using the analytical estimator proposed by Ledoit and Wolf (Ledoit and Wolf, 2004), which has been shown to perform well for this decoder, including in the context of around-ear EEG (Geirnaert et al., 2025). Training is supervised, as the attended speech envelope **y**_att_ is assumed known. Although unsupervised training strategies have been proposed that do not require attention labels (Geirnaert et al., 2021b, 2022), they are beyond the scope of this work.

At test time, given the learned decoder 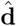, test EEG data 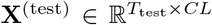 and competing speech envelopes 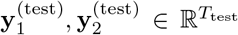, with *T*_test_ the decision window length, attentional scores *r*_1_, *r*_2_ are computed as the Pearson correlation coefficients with the reconstructed envelope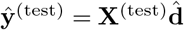 :

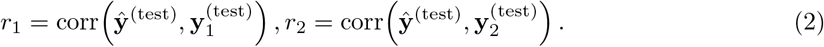

Traditionally, the speaker associated with the highest correlation is selected as the attended one. In this work, the correlation scores *r*_1_, *r*_2_ are additionally used as observations for HMM-based post-processing (Section 3.1.2).

#### 3.1.2 Hidden Markov model-based post-processing

Auditory attention exhibits temporal structure, as listeners typically maintain attention to the same speaker across successive time windows, with switches occurring relatively infrequently. Smart post-processors can exploit this temporal structure to compute a more reliable estimate of the attentional state based on the ‘observed’ attentional scores. We apply an HMM-based post-processor as proposed by Heintz et al. (2025), which has been shown to substantially improve steady-state decoding accuracy and reduce switch detection latency. Note that while no switches within a trial are present in the recorded experiment, attention switches will be simulated to enable a full evaluation of the HMM post-processing performance (see Section 3.3). While this simulation cannot fully capture real attention switch performance or behavior, it nonetheless allows a realistic evaluation of switching performance as shown in Heintz et al. (2025), where no significant difference was detected between real and simulated switch performance for the same type of algorithm. Furthermore, they extensively validated this HMM-based post-processor with real attention switches.

We employ a two-state HMM, where each state corresponds to attention to one of the two speakers, denoted by *S*_1_ and *S*_2_. The observations consist of normalized correlation score vectors **r**(*t*) =[ *r*_1_(*t*), *r*_2_(*t*) ]computed for each decision window indexed by *t* (see Eq. (2)). State probabilities P(*Ŝ*(*t*) = *S*_*j*_ | **r**_0:*t*_) are estimated using the computationally efficient forward algorithm. Note that we choose the causal version of the post-processor here, to simulate the online processing of attentional scores^1^. Emission probabilities P **r**(*t*) | *Ŝ*(*t*) = *S*_*j*_ are modeled as Gaussian distributions conditioned on the attentional state (Lopez-Gordo et al., 2025; Heintz et al., 2025).

The same hyperparameter settings are chosen as in (Heintz et al., 2025): an attention switch probability of 0.001 and a decision window length of 1 s to generate the observed correlation scores. Note that these hyperparameters are explicitly taken from Heintz et al. (2025) to guarantee honest external validation without finetuning them to this specific experiment and simulated switch settings. All correlation scores are normalized by subtracting the global mean and dividing by the global standard deviation *σ*_global_ across all (attended and unattended) correlations prior to HMM inference. Given this normalization, Gaussian emission probability distributions are chosen with standard deviations 1 and means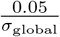 (attended) and 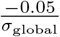 (unattended), assuming a realistic, yet again preset and untuned, average difference of 0.1 between attended and unattended correlations before normalization (Lopez-Gordo et al., 2025). For each decision window, the estimated attentional state corresponds to the state with the highest posterior probability.

### 3.2 Preprocessing

#### 3.2.1 EEG preprocessing

EEG recordings from both sensor nodes and the speech stimuli are processed on a per-trial basis. Separate EEG preprocessing is deliberately kept minimal and can be performed fully independently per node. However, it is important to remark that the data-driven stimulus decoding in Section 3.1.1 also implicitly performs and thus handles noise reduction. EEG signals are bandpass-filtered between 1–9 Hz using a zero-phase fourth-order Butterworth filter and resampled to 20 Hz. This frequency band is well established for correlation-based stimulus decoding (O’Sullivan et al., 2014; Geirnaert et al., 2021c, 2025). Finally, per-trial normalization is applied by scaling the Frobenius norm of the EEG channels to one. In case of channel selection (see Section 4.2), this normalization is performed after selection to avoid implicit overfitting.

#### 3.2.2 Speech preprocessing

As is common in sAAD studies, the clean speech envelopes are assumed available; see e.g., (Van Eyndhoven et al., 2017; Ceolini et al., 2020) for integration with speech separation. Speech envelopes are extracted following the procedure described by Biesmans et al. (2017). Speech signals are first passed through a gammatone filterbank with 19 bands spanning 50–5000 Hz, mimicking cochlear filtering. For each band, the envelope is obtained by applying a power-law compression with exponent 0.6 to the absolute value of the filtered signal. Subband envelopes are then summed, bandpass-filtered between 1–9 Hz using the same Butterworth filter as for EEG preprocessing, and resampled to 20 Hz. Finally, envelopes are z-scored on a per-trial basis.

### 3.3 Evaluation procedure

A leave-one-trial-out cross-validation (CV) scheme is used in combination with participant-specific decoders. For each participant, decoders are trained on all but one trial and tested on the held-out trial. During training, all training trials are concatenated, where the incomplete ones contribute fewer samples due to their shorter length. Although correlation-based stimulus decoding paradigms are less susceptible to such shortcut learning (Geirnaert et al., 2024), we adhere to this strict trial-based CV scheme as a best practice to avoid shortcut learning and overfitting in sAAD (Puffay et al., 2023; Rotaru et al., 2024; Ivucic et al., 2024; Yan et al., 2025). Decision window lengths are varied across *T*_test_ *∈ {*60, 30, 20, 10, 5, 2, 1} s, with each test trial segmented into non-overlapping decision windows. Raw correlation-based accuracies are primarily analyzed on 60 s windows, while 1 s windows are used for HMM post-processing.

The significance level for AAD accuracy, at *α*-level = 0.05, is computed using a random label-permutation approach, which is justified as the decision windows are non-overlapping and attentional scores computed on non-overlapping decision windows of 1 s or longer have been shown to be approximately independent (Heintz et al., 2025). Per participant and for each window length, the attention labels are 10 000 times randomly flipped with 50% probability to build a null distribution of AAD accuracies under chance. The significance level for the mean is then defined as the 95%-percentile of the distribution of averaged accuracies across participants under randomly flipped labels, while the per-participant significance levels are based on the individual null distributions. This approach automatically accounts for differences in the number of decision windows caused by incomplete trials. At the individual participant level, the consequence is that there is no global significance level; instead, each AAD accuracy is compared with the significance level for that specific participant.

To assess whether a reduced electrode configuration maintains sAAD performance (Section 4.2), a non-inferiority test is performed on the paired per-participant accuracy differences between the reduced and the original configuration on 60 s decision windows. A two-sided 90% confidence interval (CI) for the mean difference is computed using bias-corrected and accelerated bootstrapping with 10 000 resamples. The reduced configuration is considered non-inferior if the lower bound of this CI exceeds *−Δ* (corresponding to the one-sided *α*-level = 0.05), with non-inferiority marginΔ = 2 percentage points. This margin corresponds approximately to a single misclassified 60 s decision window per participant, given the average recording duration of 42 min.

For HMM evaluation, two metrics are considered: steady-state accuracy and simulated switch detection time. To enable their computation, attention switches are simulated on the correlation scores that are provided as input to the HMM. First, all 1 s correlation scores are concatenated across trials and reordered such that the first score always corresponds to the attended speaker. Attention switches are then generated randomly across the entire recording at an average rate of 2.5 switches per 10 min, with a minimum inter-switch interval of 2 min. A switch is implemented by swapping the correlation scores, effectively reversing the attention labels. Switch detection time is defined as the latency required for the posterior probability of the newly attended speaker to exceed that of the previously attended speaker. Given the two-speaker setup, in practice this boils down to the time needed for the newly attended speaker to reach *≥* 50% attended probability. Steady-state accuracy is computed based on all posterior probabilities across all decision windows, excluding those within the switch transition periods that determine the switch detection time to reduce cross-contamination between both metrics. This switch simulation procedure is performed 100 times to reliably estimate the average steady-state accuracy and switch detection time. A similar switch simulation procedure for the same type of algorithm has been validated in Heintz et al. (2025), where no significant difference was detected between real and simulated switch performance.

## 4 Results

This section presents the experimental results obtained with the proposed wireless EEG sensor network. We first compare the performance of combining sensor nodes from both ears with single-ear configurations (Section 4.1). Next, we analyze bipolar channel selection strategies to identify practically optimal electrode configurations within the WESN setup (Section 4.2). Finally, we evaluate the impact of HMM post-processing (Section 4.3).

### 4.1 Comparison both ears combined versus single-ear configurations

We first investigate whether combining sensor nodes from both ears improves sAAD performance compared to single-ear configurations. Figure 2a shows the average sAAD accuracy as a function of decision window length, based on the raw neural tracking correlations in Eq. (2). Figure 2b reports per-participant accuracies for 60 s decision windows.

**Figure 2.**
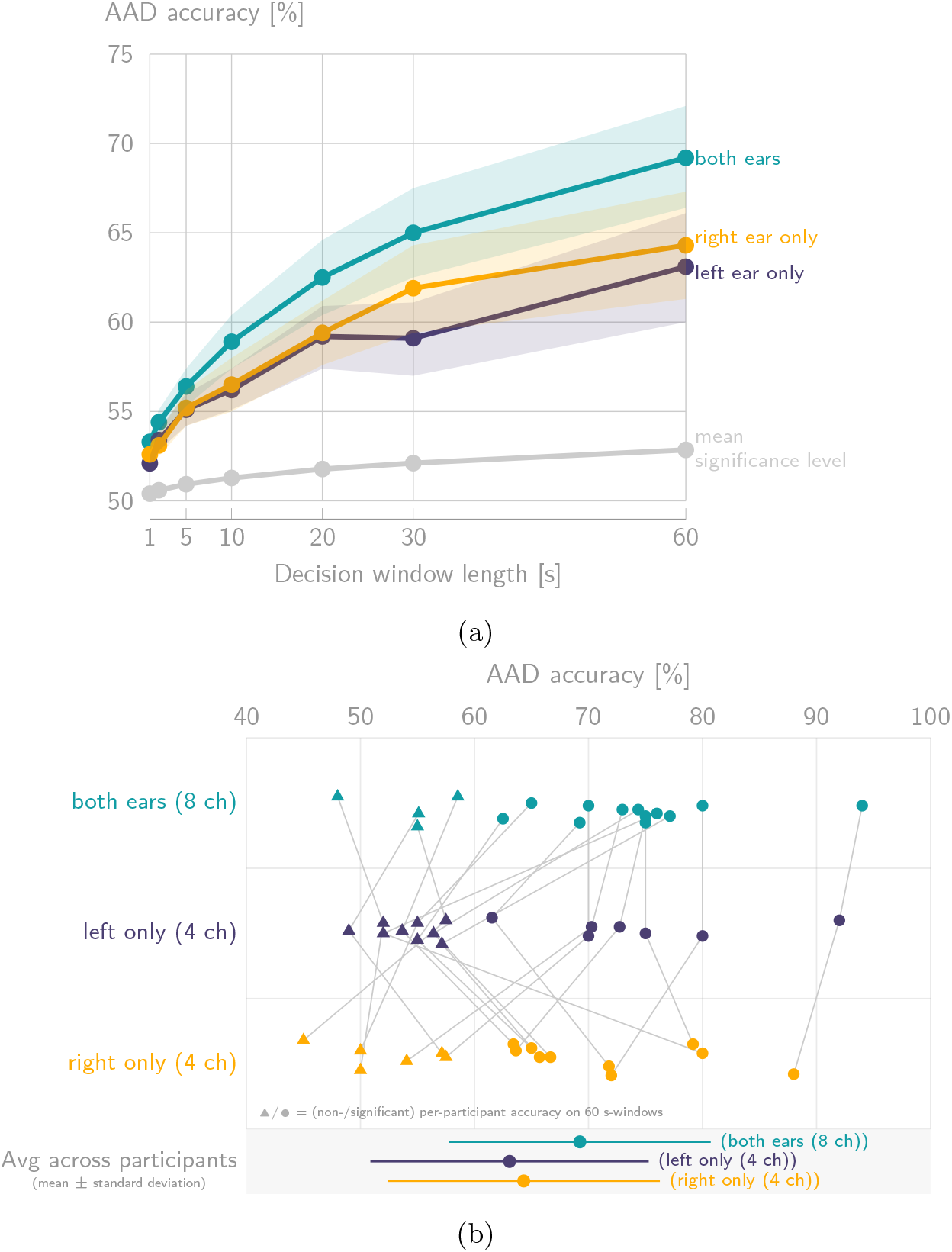
**(a)** The average sAAD performance curves (mean *±* standard error across the 16 participants; mean significance level computed based on random label-permutation, seeSection 3.3) based on raw neural tracking correlations show that using wireless EEG sensor nodes at both ears simultaneously substantially outperforms single-ear configurations. **(b)** The per-participant sAAD accuracies for 60 s decision windows. Gray lines connect results from the same participant; participants with non-significant AAD accuracies (based on per-participant significance levels using random label-permutation, see Section 3.3) are indicated using a triangle.

When using all available channels, combining both ears yields an average accuracy of 69.24%, compared to 63.07% for the left ear only and 64.32% for the right ear only (60 s windows). All three settings exhibit significant decoding accuracy on average, as seen in Figure 2a. At the individual level for 60 s windows, 12*/*16 participants exhibit significant AAD accuracy for both ears, 7*/*16 for the left ear only, and 10*/*16 for the right ear only (Figure 2b). Using a paired Wilcoxon signed-rank test with Bonferroni–Holm correction (*α* = 0.05, two comparisons), performance with both ears (all channels) is significantly higher than with the left ear only (uncorrected *p* = 0.0061, corrected *p* = 0.0122; *W* = 8, rank-biserial *r* = 0.727, *N* = 16) and the right ear only (uncorrected *p* = 0.0386, corrected *p* = 0.0386; *W* = 28, rank-biserial *r* = 0.517, *N* = 16).

To investigate how combining both ears compares to single-ear configurations across a varying number of channels, Figure 3 illustrates the effect of limiting the number of channels per ear on the sAAD accuracy based on the raw correlations in Eq. (2). Across all channel combinations (taken from the original set of four channels per ear) and participants, combining both ears consistently outperforms single-ear configurations by approximately 5 percentage points, regardless of the number of channels per ear. For the combined-ear configuration, selected channels are always matched across both ears.

**Figure 3.**
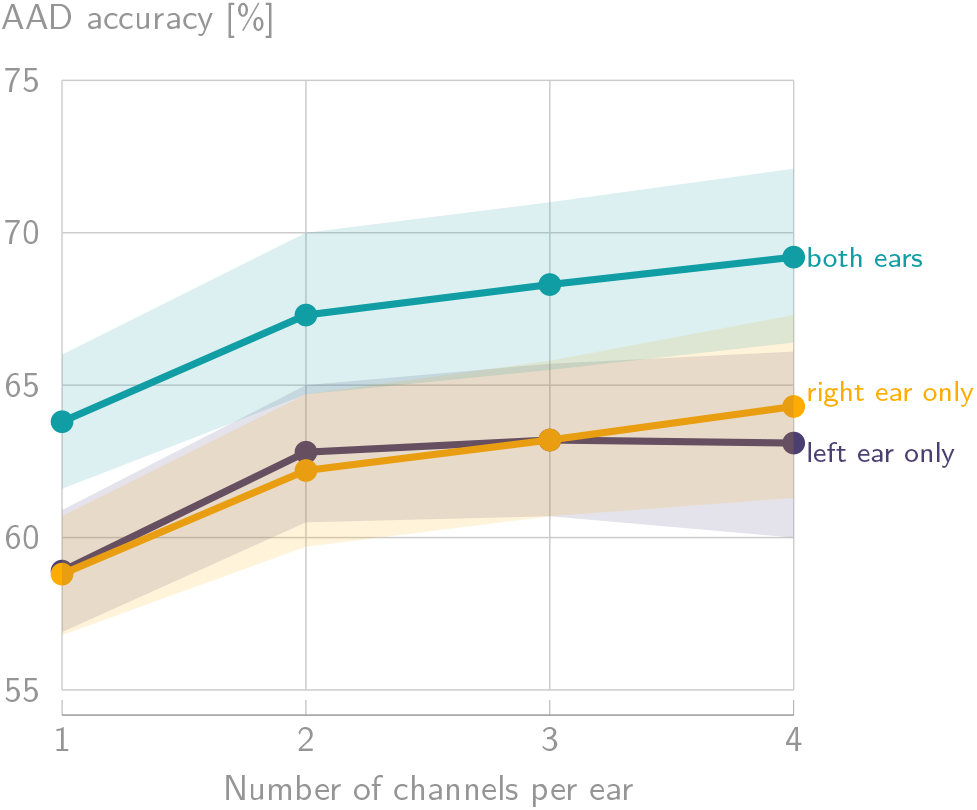
The average sAAD accuracy (mean *±* standard error across the 16 participants and all channel combinations) for 60 s decision windows as a function of the number of channels per ear shows a consistent 5 percentage point difference in accuracy between combining both ears and single-ear configurations.

### 4.2 Channel selection using bipolar configurations

Each sensor node provides five electrodes: four around-ear electrodes (E1-4) and one reference electrode (REF) (see Figure 1a). In the previous analyses, four monopolar channels per ear were used by referencing each around-ear electrode to the local reference electrode. However, alternative bipolar configurations may yield more informative dipoles. We therefore perform a channel selection analysis considering all possible bipolar channels.

Per ear, the five electrodes allow for ten possible bipolar channels. For a given number of channels per ear (ranging from one to four), channel selection is performed using a leave-one-participant-out procedure combined with participant-specific decoding as before. Specifically, for each left-out participant, optimal channel combinations are selected based on the average performance across the remaining participants using leave-one-trial-out CV. These selected channels are then evaluated on the left-out participant, again using participant-specific leave-one-trial-out CV. When combining both ears, channel selection is constrained to use identical channel configurations for the left and right ears. Additionally, channel combinations are restricted to use at most one additional electrode beyond the number of channels, reflecting practical hardware constraints.

Figure 4a shows the resulting average sAAD accuracies for 60 s decision windows. Bipolar channel selection mostly yields (slightly) higher accuracies compared to the original monopolar configurations. Compared to the original configuration of four monopolar channels per ear, the three-channel bipolar configuration (using four electrodes per ear and combining both ears) yields a mean paired difference of +0.89 percentage points (90% CI = [*−* 1.25, 2.58]). According to the non-inferiority test described in Section 3.3 with marginΔ = 2 percentage points, this three-channel bipolar configuration is thus non-inferior to the original four-channel configuration.

**Figure 4.**
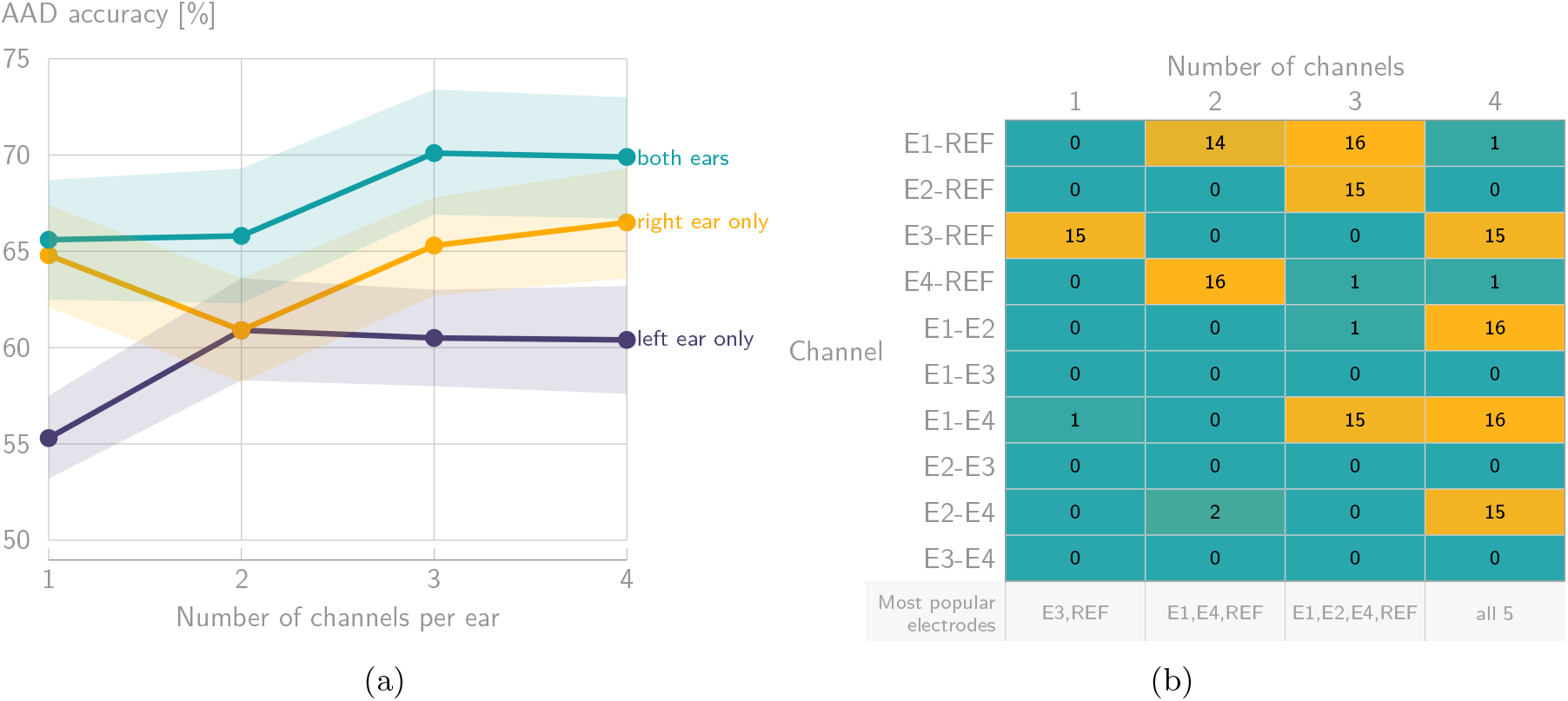
**(a)** The average sAAD accuracy (mean *±* standard error across the 16 participants) on 60 s decision windows using a bipolar channel selection as a function of the number of channels per ear shows that three channels using four electrodes per ear are sufficient to reach around 70%. **(b)** The heatmap of selected bipolar channels across participants for the combined-ear configuration indicates the most popular channels and required electrodes to be included in a practical WESN configuration.

Figure 4b summarizes the frequency with which specific bipolar channels are selected across participants when combining both ears. Based on the most frequently selected channels, a uniform bipolar configuration using three channels per ear and four electrodes (channels E1–REF, E2–REF, and E1–E4) achieves an average accuracy of 70.63% prior to HMM post-processing. Note that, as this configuration is derived from the channel selection across all participants, this accuracy is slightly optimistic compared to the unbiased estimate of 70.13% from the leave-one-participant-out procedure.

### 4.3 HMM post-processing results

Finally, we evaluate the impact of HMM-based post-processing, which exploits the temporal structure of attentional processes as explained in Section 3.1.2. Attention switches were simulated as described in Section 3.3. Figure 5 shows per-participant steady-state accuracies and average switch detection times for the combined-ear configuration, clearly showing how better-performing participants yield both improved steady-state accuracy and lower average switch detection times.

**Figure 5.**
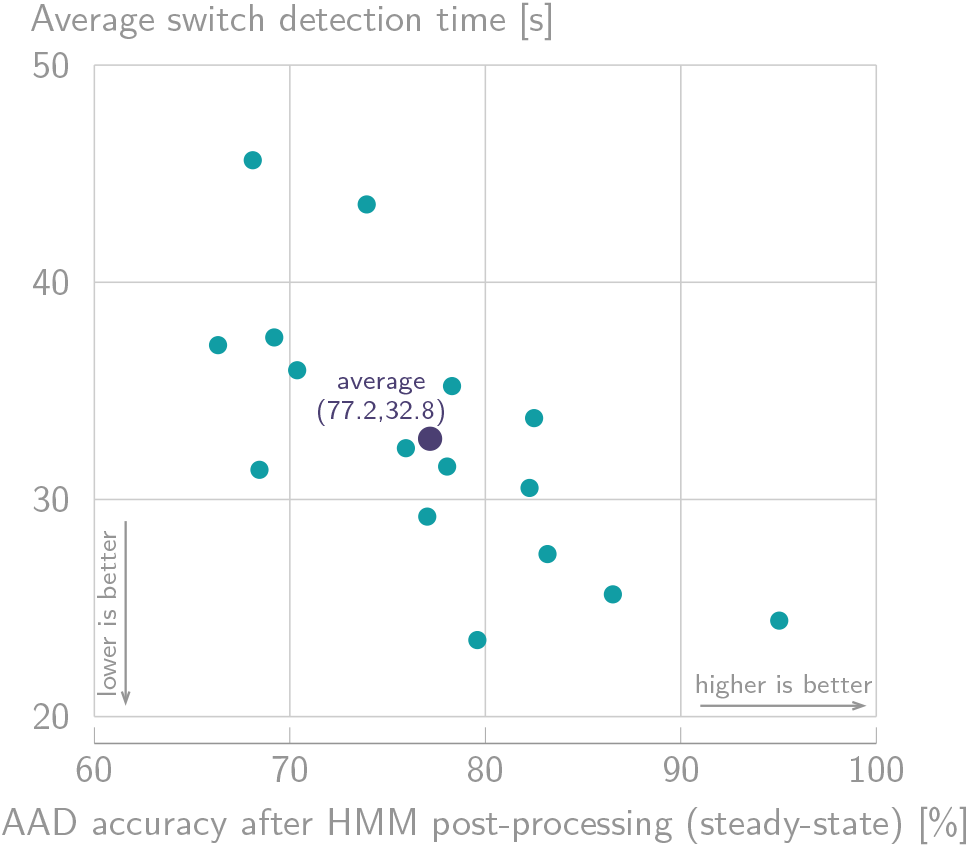
The per-participant steady-state sAAD accuracies and average simulated switch detection times after HMM post-processing for the combined-ear configuration show a clear improvement compared to the raw sAAD accuracies shown in Figure 2b.

HMM post-processing increases the average sAAD accuracy using all four channels from both ears from 69.24% (60 s decision windows) to 77.17%, while achieving a mean switch detection time of 32.79 s. For single-ear configurations, HMM post-processing similarly improves performance, yielding average accuracies of 74.10% (left ear, 35.41 s switch detection time) and 75.03% (right ear, 32.19 s switch detection time). The trade-off between steady-state accuracy and switch detection time can be further tuned by adjusting the HMM transition probabilities, yet this is beyond the scope of this paper. Lastly, the optimal bipolar configuration from Section 4.2, using three channels per ear and four electrodes, reaches an average accuracy of 77.08% after HMM post-processing, with an average switch detection time of 33.40 s.

## 5 Discussion

Based on the results presented in Section 4, three main discussion points can be identified and are elaborated below.

### 5.1 Selective auditory attention can be decoded using synchronized and galvanically separated EEG sensor nodes around the ear

First, the successful and significant sAAD performance obtained when combining two wireless EEG sensor nodes positioned around both ears (see, e.g., Figure 2a) demonstrates the practical viability of the developed WESN platform (Ding et al., 2025b), including its synchronization protocol (Section 2.1.3). These results confirm that synchronized, battery-powered, and galvanically separated miniature EEG sensor nodes can be reliably employed in a practical neuro-steered hearing device setting, yet still under controlled laboratory conditions.

Second, the decoding performance obtained using the miniature around-ear WESN setup is highly comparable to the performance reported in previous around-ear-based studies, which typically relied on a single physically connected EEG grid with a shared reference electrode and achieved approximately 70% decoding accuracy on 60 s decision windows (Mirkovic et al., 2016; Holtze et al., 2022; Geirnaert et al., 2025). In the present work, galvanic isolation between the two nodes prevents the measurement of long-distance cross-ear potential differences. Nevertheless, this constraint does not appear to degrade sAAD performance. Although this comparison is made across studies, and thus across participants, stimuli, and recording setups, it is corroborated by earlier findings by Geirnaert et al. (2025), where galvanic isolation was emulated in around-ear EEG within the same participants and recordings, and no performance loss was observed. Taken together, these results suggest that little additional neural information for effective sAAD can be captured from cross-ear potential differences, but rather that most information is decoded from same-ear potentials.

Third, the HMM-based post-processing results reported in Section 4.3 provide a realistic performance benchmark for sAAD using a practical WESN. A steady-state decoding accuracy of 77.17% was achieved, with an average simulated switch detection time of 32.79 s (based on idealized, instantaneous simulated switches; future work should include validation on real attention switches). While acceptable performance thresholds for real-world hearing device users remain unclear, these values are likely insufficient for a fully satisfactory user experience aimed at substantially improving speech intelligibility. Importantly, when HMM output probabilities are directly mapped to per-speaker gain control, the reported switch detection time corresponds to the point in time at which the relative playback levels of both speakers become equal following an attention switch. However, the perceptual gain adaptation towards the newly attended speaker begins much earlier, as the HMM relative output probabilities evolve smoothly over time (Heintz et al., 2025). As a result, the listener may experience a gradual enhancement of the attended speech well before the formal switch detection point. Moreover, while the obtained performance metrics may be insufficient for highly dynamic listening scenarios, many everyday listening scenarios are characterized by more slowly varying attentional dynamics (Hjortkjær et al., 2025), also enabled by, for example, grouping speech streams in conversations (Van de Ryck et al., 2025). Within this context, the presented results provide a baseline for what can currently be achieved if implemented in neuro-steered hearing devices, against which future methodological and hardware improvements can be evaluated.

### 5.2 Combining sensor nodes around both ears outperforms single-ear configurations primarily through increased robustness

The results consistently demonstrate a performance advantage when combining EEG sensor nodes at both ears compared to using single-ear configurations (Figures 2 and 3), with statistically significant differences observed at the group level and more participants exhibiting significant accuracies at the individual level. Beyond confirming the successful synchronization of the two wireless nodes, these findings indicate that a binaural EEG configuration is beneficial for achieving higher and more reliable sAAD performance.

Importantly, further analysis suggests that this benefit is driven primarily by increased robustness rather than by consistently exploitable spatially complementary neural information across ears. Specifically, when considering the post-hoc best-performing single-ear configuration for each participant across all trials, an average accuracy of 68.65% is obtained on 60 s decision windows, which is not significantly different from the 69.24% achieved by the combined-ear configuration (paired Wilcoxon signed-rank test; *p* = 0.8911). Moreover, for approximately half of the participants, the left ear yielded the best single-ear performance, while for the other half, the right ear performed best (Figure 2b). This inter-individual variability indicates that combining both ears mainly provides redundancy that mitigates ear-specific performance differences, leading to a more robust decoding performance across users. While complementary information across ears cannot be entirely ruled out, the present results suggest that robustness is the dominant factor underlying the observed performance gains. An important question for future research is whether, within individual participants, the better-performing ear remains consistent across recording sessions or instead reflects coincidental, session-specific factors, such as the specific experimental setup and conditions, electrode placement, or electrode impedances. Nonetheless, in practice, the best-performing single-ear configuration is in any case not known beforehand, warranting the use of the more robust both-ear configuration.

### 5.3 Three channels using four electrodes per ear are sufficient for robust sAAD

The channel selection analysis presented in Section 4.2 demonstrates that three bipolar channels derived from four electrodes per ear are non-inferior to the original four channels using five electrodes per ear. Using the post-hoc selected fixed bipolar configuration consisting of channels E1–REF, E2– REF, and E1–E4 (across both ears), and thus relying on electrodes E1, E2, E4, and REF only (see Figure 1a), an average decoding accuracy of 70.63% before and 77.08% after HMM post-processing (with an average switch detection time of 33.40 s) was obtained.

These results indicate that a reduced and participant-independent electrode layout can be employed without a meaningful loss in decoding performance, which is highly relevant for the design of wearable, concealable, and user-friendly neuro-steered hearing devices. In contrast, further reductions in the number of electrodes result in noticeable performance degradation (Figure 4a). Overall, this analysis provides concrete design guidelines for integrating around-ear EEG into a practical WESN-based neuro-steered hearing device, balancing decoding performance against wearability and system complexity. In future work, the final three-channel configuration relying on electrodes E1, E2, E4, and REF should be validated on an independent cohort. Furthermore, additional mini-EEGs that do not compromise wearability and unobtrusiveness, such as in EEG glasses, could be seamlessly added through this WESN platform to further improve decoding performance, while dry electrodes and out-of-the-lab conditions should also be tested.

## 6 Conclusion

In this work, we established a proof of concept of a wireless EEG sensor network (WESN) consisting of two synchronized and galvanically separated around-ear EEG sensor nodes for selective auditory attention decoding in neuro-steered hearing devices. Using a traditional correlation-based stimulus decoding algorithm, an average decoding accuracy of 69.24% was achieved on 60 s decision windows, which is in line with the performance reported in the literature for physically connected around-ear EEG setups with a shared reference.

We further demonstrated that combining sensor nodes at both ears consistently outperforms single-ear configurations, and hypothesized that this is primarily driven by providing redundancy that increases robustness across users rather than by exploiting additional cross-ear neural information. Using HMM-based post-processing, a realistic performance benchmark for sAAD with a practical WESN was established, yielding a steady-state accuracy of 77.17% with an average simulated switch detection time of 32.79 s. This benchmark provides a meaningful reference point for future practical implementations of neuro-steered hearing devices based on wireless EEG. Finally, through a bipolar channel selection analysis, we showed that the number of electrodes per ear can be reduced from five to four (from four to three channels) using a fixed and participant-independent configuration, without a meaningful loss in decoding performance.

Overall, this work demonstrates that selective auditory attention decoding is feasible using a fully wireless, synchronized, and galvanically isolated EEG sensor network with unobtrusive around-ear EEG, representing a considerable step towards practical and wearable neuro-steered hearing devices.

## Acknowledgments

ChatGPT (version 5.2; OpenAI) and Claude Sonnet (version 4.6; Anthropic) were used as a writing support for language editing and grammar checking. Claude Sonnet (version 4; Anthropic) was used as a coding assistant for software development and implementation support. All algorithms, analyses, and interpretations were designed, verified, and validated by the authors, who take full responsibility for the results. All authors have reviewed and approved the final content of the manuscript and take full responsibility for the paper’s scientific integrity.

The authors thank Joachim Rivera Cea for providing the trigger implementation.

## Funding

This research is funded by the Research Foundation - Flanders (FWO) project No G081722N, a junior and senior postdoctoral fellowship fundamental research of the FWO (for S. Geirnaert, No. 1242524N and 1211827N), the European Research Council (ERC) under the European Union’s research and innovation program (grant agreement No 802895 and No 101138304), Internal Funds KU Leuven (project IDN/23/006), and the Flemish Government (AI Research Program). Views and opinions expressed are however those of the author(s) only and do not necessarily reflect those of the European Union or the granting authorities. Neither the European Union nor the granting authorities can be held responsible for them.

## Author contributions

*Simon Geirnaert* : Conceptualization, Data curation, Formal analysis, Funding Acquisition, Investigation, Methodology, Software, Visualization, Writing - original draft, Writing - review & editing. *Ruochen Ding* : Conceptualization, Data curation, Investigation, Resources, Writing - original draft, Writing - review & editing. *Alexander Bertrand* : Conceptualization, Formal analysis, Funding acquisition, Project administration, Resources, Supervision, Writing - review & editing.

## Data availability

The full dataset will be made publicly available upon acceptance.

## Footnotes

1 As this work focuses on offline evaluation, the overall pipeline is still non-causal, for example, due to the preprocessing in Section 3.2.1. The simulation of online processing only applies to the HMM inference.

## Notes

### Competing Interest Statement

The authors have declared no competing interest.

### Summary of Updates

Added statistical non-inferiority test to show that channel-reduced setup is not worse than full setup + rephrasings and some extra literature to back up claims.

